# Microbiota assembly of specific pathogen-free neonatal mice

**DOI:** 10.1101/2025.01.14.633035

**Authors:** Elizabeth A. Kennedy, James S. Weagley, Andrew H. Kim, Avan Antia, Anna L. DeVeaux, Megan T. Baldridge

## Abstract

**Background:** Neonatal mice are frequently used to model diseases that affect human infants. Microbial community composition has been shown to impact disease progression in these models. Despite this, the maturation of the early-life murine microbiome has not been well-characterized. We address this gap by characterizing the assembly of the bacterial microbiota of C57BL/6 and BALB/c litters from birth to adulthood across multiple independent litters.

**Results:** The fecal microbiome of young pups is simple, dominated by only a few pioneering bacterial taxa. These taxa are present at low levels in the microbiota of multiple maternal body sites, precluding a clear identification of maternal source. The pup microbiota begins diversifying after fourteen days, coinciding with the beginning of coprophagy and the consumption of solid foods. Pup stool bacterial community composition and diversity are not significantly different from dams from day 21 onwards. Short-read shotgun sequencing-based metagenomic profiling of young pups enabled the assembly of metagenome-assembled genomes for strain-level analysis of these pioneer *Ligilactobacillus, Streptococcus,* and *Proteus* species.

**Conclusions:** Assembly of the murine microbiome occurs over the first weeks of postnatal life and is largely complete by day 21. This detailed view of bacterial community development across multiple commonly employed mouse strains informs experimental design, allowing researchers to better target interventions before, during, or after the maturation of the bacterial microbiota. The source of pioneer bacterial strains appears heterogeneous, as the most abundant taxa identified in young pup stool were found at low levels across multiple maternal body sites, suggesting diverse routes for seeding of the murine microbiome.

## Background

Newborn human infants gradually acquire a diverse set of microbes starting at the time of delivery [1]. The gut microbiota matures over the first years of life and is shaped by the mode of delivery, nutrition, and exposure to antibiotics [2–4]. Birth mode is the main factor known to differentiate the gut microbiome immediately after birth, with vaginally born infants exhibiting microbiomes enriched for *Lactobacilli*. In contrast, infants delivered by Caesarean section are colonized by genera such as *Staphylococcus* and *Propionibacterium* [5–7]. Delivery mode continues to affect microbial populations for the first few years of life [3]. The diet also influences the progression of early life gut microbiome composition. Breast-fed infants have more *Bifidobacterium* and *Lactobacillus* in their gut microbiota compared to formula-fed infants [2, 8]. A milk-based diet selects for microbes that can digest milk oligosaccharides, and the introduction of solid foods induces a shift towards microbes that can digest a wider set of macromolecules such as *Bacteroides* and *Clostridia* [2, 9, 10]. Gut microbes change as infants are exposed to a broader variety of environments, such as exposure to other family members, pets [11], and daycare [12], but ultimately achieve an adult-like configuration by the age of three [13].

Neonatal mice are widely used as models to study infectious and inflammatory conditions associated with human infants. Early-life bacterial communities affect the course of various diseases, including rotavirus [14], *Cryptosporidium* [15], and *Salmonella* infections [16], as well as necrotizing enterocolitis [17]. Previous studies have found that the mouse microbiome immediately after birth has taxa overlapping with the maternal vaginal microbiome [18], but the gut microbiota composition rapidly shifts over the first 24 hours of life, likely representing pioneer microbes that are unable to stably colonize the neonatal gut [19]. At weaning, the pup’s microbial composition is similar to the fecal microbiota of the dam, as the pups become coprophagic [20, 21]. The source of the full community of pre-weaning neonatal gut microbes remains somewhat unclear, although exposure to the microbiota of other maternal body sites, such as the skin, may contribute to neonatal gut microbe populations in Caesarean-section delivered neonates [22].

To date, the early-life transitions of the murine enteric microbiota through weaning and into adulthood have not been well-profiled. Without the ability to predict the approximate complexity or conformation of bacterial communities likely to be present at a given pup age, optimal experimental design for challenges administered pre-weaning is encumbered. Here, we characterized the pup microbiota of specific pathogen-free (SPF) litters throughout the first three weeks of life as well as into adulthood and observed predominantly *Ligilactobacillus* with some contributions from *Streptococcus* and *Proteus* as dominant early-life microbes in the SPF setting. After approximately two weeks of life, the mouse microbiota transitioned from a very simple community to a more complex, adult-like community, associated with the pup’s dietary transition from breastmilk to chow along with coprophagic behavior. This longitudinal profiling of the early-life microbiota provides an important window into the timing of microbiota maturation as well as the taxonomic identities of bacteria associated with these transitions. Additionally, short-read shotgun sequencing-based metagenomic profiling enabled the analysis of strain-level variation of the early murine microbiome. Together, these analyses provide a comprehensive view of bacterial microbiome development in the mouse gastrointestinal tract.

## Methods

### Mice

Pregnant dams (E13-E16 on arrival) were purchased from Charles River (BALB/c, strain #028; C57BL/6, strain #027) and gave birth shortly after arrival at Washington University in Saint Louis 3-7 days later. Dams were housed under specific pathogen-free conditions with autoclaved standard chow pellets and water provided *ad libitum*. Pups were weaned and separated into cages by sex at postnatal day 21, with no more than 5 mice per cage. Animal protocols 20190162 and 22-0140 were approved by the Washington University Animal Studies Committee.

### Collection of fecal samples from neonates and dams

Samples were collected from neonates beginning shortly after birth until weaning at postnatal day (P)21, then weekly until 6 weeks old. Fecal samples were collected from dams at the first sampling of the pups (P4), days later (P7/8), and at weaning (P21). Fecal samples were harvested into 2 mL tubes (Sarstedt, Nümbrecht, Germany) with 1-mm-diameter zirconia/silica beads (Biospec, Bartlesville, OK) and stored at −80°C until processing.

### Collection of samples from maternal body sites

Face (both cheeks) and ventral samples were collected by vigorous swabbing of maternal skin with a sterile swab soaked in lysis buffer (200 mM NaCl, 200 mM Tris, 20 mM EDTA). Swabs were spun into an Eppendorf tube using Lyse&Spin collection tubes (Qiagen, Hilden, Germany). Maternal oral and vaginal samples were collected by repeated lavage with PBS (50 μL PBS, 4x washes per site). ‘Sample collection’ negative control swabs and PBS samples were collected and processed alongside each set of maternal samples.

### 16S rRNA gene amplicon sequencing of feces

DNA was extracted from fecal pellets using phenol:chloroform extraction followed by clean-up using the DNeasy Blood and Tissue Kit (Qiagen, Hilden, Germany). Amplicons were generated using barcoded PCR primers targeting the V4 region of the 16S rRNA gene as described previously, with 26 cycles of PCR, and purified using Agencourt Ampure XP beads [23]. Amplicon sequencing of the 16S rRNA gene V4 gene region of the fecal samples generated 8.99 × 10^6^ sequences with a median of 22,153 reads per sample.

### 16S rRNA gene amplicon sequencing of maternal body sites

Maternal body site samples were processed in the same manner as fecal pellets, except 30 cycles of PCR were run to amplify the V4 region. Maternal body site samples were pooled and sequenced separately from fecal samples to ensure adequate sequencing coverage for these low-biomass sites. Multiplex sequencing was performed on an Illumina MiSeq instrument (bi-directional 250 nucleotide reads) generating 405,014 sequences with a median of 5,410 reads per sample.

### Quality control of sequencing data from maternal body sites

Maternal body sites harbored substantially lower bacterial biomass than fecal samples and required more PCR amplification before sequencing, increasing the potential for contamination during sample collection and processing. A ‘sample processing’ negative control was included in addition to the previously mentioned ‘sample collection’ negative controls gathered during maternal sampling. These control samples were sequenced and analyzed in parallel with body site samples. Read counts for all controls were on average lower than samples, but many samples from the low biomass sites had read counts equivalent to or below controls (**Fig S1**). Face swabs *p*=0.0144), but not ventral swabs (*p*=0.0623), had significantly higher read counts than negative control swab samples. Vaginal and oral samples had significantly higher read counts than PBS wash controls (*p*=0.0465 and *p*=0.0024, respectively). Taxonomic analysis of the maternal microbiome was constrained to samples with greater than 1,500 reads, a cutoff based on read counts of negative controls.

### Quantitative PCR of the 16S rRNA gene

SYBR green-based quantitative PCR of the 16S rRNA gene was performed in duplicate using 515F (5’-GTGCCAGCMGCCGCGGTAA-3’) and 805R (5’-GACTACCAGGGTATCTAATCC-3’) primers on phenol:chloroform-extracted DNA from mouse fecal pellets. Absolute copies were quantified based on a standard curve.

### Processing and analysis of 16S rRNA gene amplicon sequencing data

Sequences were processed using mothur’s MiSeq standard operating procedure [24]. Raw fastq files were demultiplexed and quality filtered. Chimeras were identified using mothur’s implementation of VSEARCH[25] and removed. Sequences were classified using the RDP reference taxonomy database (version 16) and sequences identified as mitochondria or chloroplast were removed. Linear discriminant analysis Effect Size (LEfSe) analysis was used to determine discriminatory taxa between age groups [26]; an LDA effect size of 4.0 was used as the cutoff for reported taxa.

### Short-read shotgun sequencing of fecal gDNA

Seven sequencing libraries were generated from gDNA previously extracted from pup feces using the Nextera DNA Library Prep Kit (Illumina, San Diego, CA) and barcoded primers. Samples were selected based on the abundance of bacterial taxa identified via amplicon sequencing; selected samples were enriched for dominant early-life taxa from BALB/c and C57BL/6 pups across multiple litters. Libraries were sequenced on an Illumina MiSeq instrument [bi-directional 150 nucleotide reads; 3.45 × 10^6^ ± 1.53 × 10^6^ reads/sample (minimum of 1.20 × 10^6^ reads)]. Sequences were adapter and quality trimmed using TrimGalore (v. 0.6.8 dev) [27] and cutadapt (v. 2.10) [28]. Quality control was performed using FastQC (v. 0.11.9) [29].

### Generation and classification of metagenome-assembled genomes

Metagenome-Assembled Genomes (MAGs) were generated using MEGAHIT (v. 1.2.9) [30] and binned using anvi’o interactive software (v. 7.1) [31]. Bins were generated based on both sequence composition and coverage statistics – contigs arising from the same genome have similar sequence compositions and their coverage covaries across samples based on organismal abundance. Taxonomy was assigned to reads using Centrifuge (v. 1.0.4) against the NCBI nucleotide reference database [32]. Reads were mapped to contigs using bowtie (v. 2.3.5) [32], allowing taxonomy to be assigned to contigs and bins. Genome completeness and redundancy were calculated for each bin within anvi’o [31].

### Annotating viruses and plasmids in metagenomic assemblies

VirSorter2 (v. 2.2.4) was used to identify viruses in all contigs generated in the previously described metagenomic assemblies. PhaTYP [33] and PhaGCN [34] were used to generate lifestyle and taxonomic predictions, respectively, for the identified viruses. Mmseqs2 was subsequently used to search non-redundant nucleotide sequences from the gut phage database (GPD) and identified 22 phages with E-value ≤ 4.09 x 10^-4^ [35, 36]. Metadata associated with these hits were used as a secondary source of information regarding the taxonomic identity of phages and their bacterial hosts. Plasmer (downloaded September 20, 2023) [37, 38], a random forest classifier trained on k-mer frequencies and other genomic features, was employed to identify contigs of plasmid origin.

## Results

### The simple early-life mouse microbiota begins diversifying around postnatal day 15

To characterize the maturation of the mouse microbiome throughout development, we obtained pregnant C57BL/6 and BALB/c dams from Charles River and characterized the bacterial composition of their litters from neonates to adulthood using amplicon sequencing of the v4 region of the16S rRNA gene. Taxonomic classification of fecal samples collected from pups revealed age- and litter-specific differences in the gut microbiota (**Fig 1A, Fig S2**). Early in life (P4-P14), the neonatal microbiota was generally dominated by *Ligilactobacillus* with a small proportion of *Streptococcus*, with one litter exhibiting a robust representation of *Proteus*. Around P15, those dominant taxa decreased and Bacteroidetes prevalence increased. LEfSe analysis, performed to identify differentially abundant taxa before and after P14, indicated that Firmicutes, primarily *Ligilactobacillus* and *Streptococcus*, were discriminatory for P10-P14 samples (**Fig S3)**. In contrast, Bacteroidetes, primarily *Muribaculaceae* and *Bacteroides*, and Clostridia, primarily *Lachnospiraceae*, were discriminatory for P15-P20 samples. *Lactobacillus*, which is closely related to *Ligilactobacillus*, was differentially abundant in older pups (**Fig S3**).

**Figure 1:**
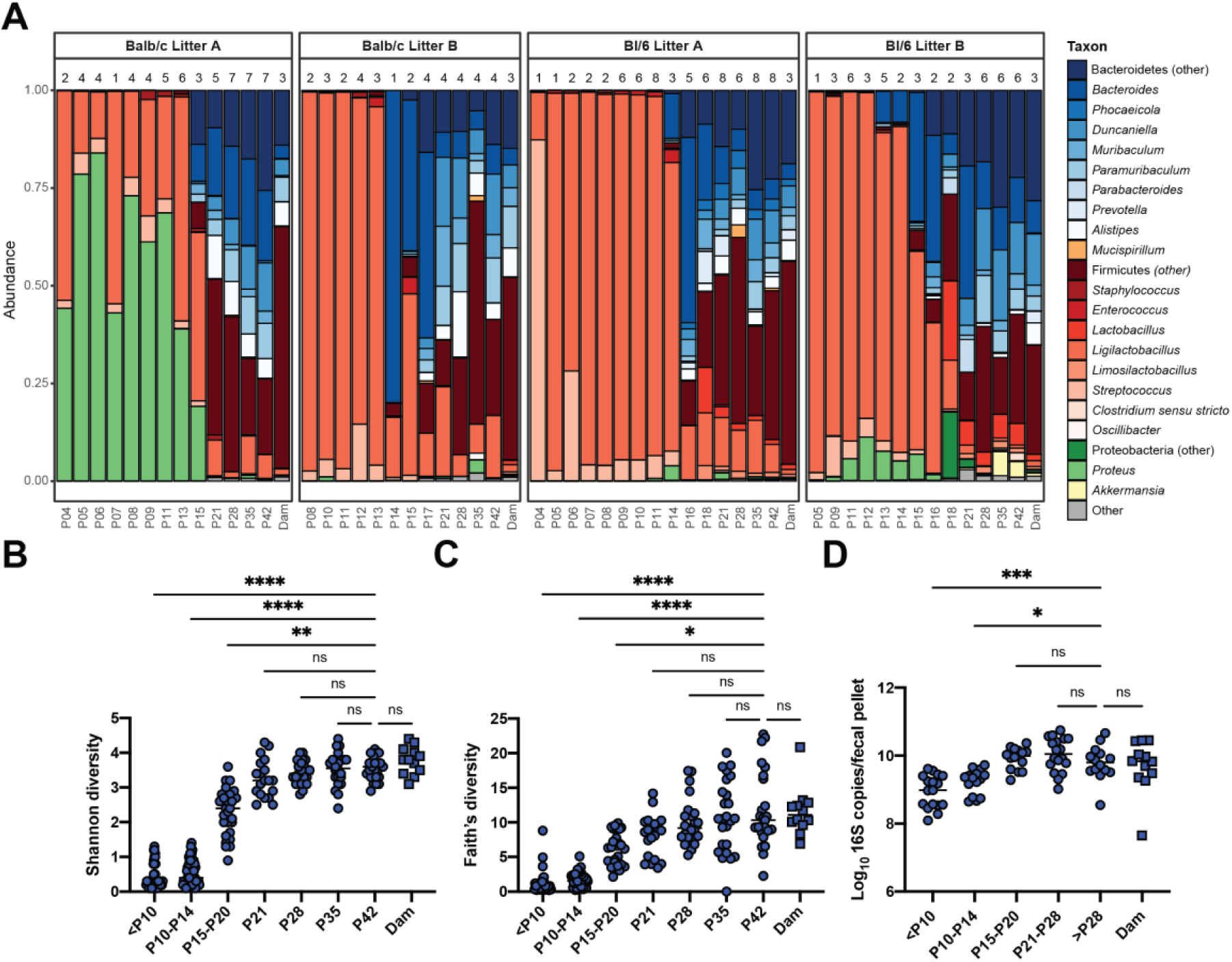
The early-life fecal microbiota begins diversifying at postnatal day 15. **(A)** Taxonomic classification of stool samples collected from neonates sequenced at the 16S rRNA gene V4 region. Genera represented at greater than 5% abundance in at least one sample are shown. Each bar represents the average abundance of taxa from all samples collected from pups from the litter for a given time point; the n above each bar indicates the number of samples collected for that time point. Color families show phyla-level assignments – blue for Bacteroidetes, orange for Deferribacteres, red for Firmicutes, green for Proteobacteria, and yellow for Verrucomicrobia; grey includes phyla present at less than 5% abundance in all samples. ( **B)** Shannon diversity calculated based on operational taxonomic unit (OTU) clustering. **(C)** Faith’s phylogenetic diversity was calculated based on the phylogeny of amplicon sequence variants (ASVs). **(D)** 16S rRNA gene copies per fecal pellet, detected by qPCR. Medians are indicated by a horizontal line. Results were compared by the Kruskal-Wallis test with Dunn’s test for multiple comparisons. * *p* < 0.05, *\*\* p* < 0.01, \**\*\* p* < 0.001, \*\**\*\* p* < 0.0001, ns = not significant; *n* =12 – 57, representing samples from mice combined across four litters from two genotypes as in **A** within indicated age ranges.

Microbiota alpha diversity was calculated using the Shannon diversity index of sequences clustered into operational taxonomic units (OTUs) (**Fig 1B, Fig S4A**). Diversity was low until P14, then increased until the age of weaning and remained stable through sampling at 6 weeks, at which time it was comparable to the diversity of the dam’s fecal microbiota. The average Shannon diversity of P10-P14 samples (mean Shannon diversity ± SD = 0.57 ± 0.32, n=56) was significantly lower than P15-P20 samples (mean Shannon diversity ± SD 2.35 ± 0.70, n=27; Mann-Whitney p- value < 0.0001). Analysis of Faith’s phylogenetic diversity, calculated based on the phylogeny of amplicon sequence variants (ASVs), showed low initial diversity which increased at P15 through weaning and then stabilized (**Fig 1C, S4B**). Measurement of absolute 16S rRNA gene levels in stool samples by quantitative PCR indicated that absolute bacterial loads were lower in early life compared to adult samples (**Fig 1D, S4C**). These data support that the early-life bacterial microbiota has relatively low diversity and begins diversifying with associated compositional shifts around 15 days of life before stabilizing around the time of weaning.

### Pup fecal bacterial community structure shifts with age and shares features with dam fecal microbiota

We assessed bacterial community structure by clustering OTUs and comparing longitudinal samples within a litter based on the Yue-Clayton theta similarity index, which considers both OTU presence/absence and relative abundance. Age was a key driver of community structure changes as revealed by principal coordinate analysis (PCoA) for each litter, with samples from pups younger than P15 (P ≤ 14) clustering together before shifting to an intermediate configuration (P15-P21) and finally to an adult-like community (**Fig 2A**). The P21 pup stool bacterial community was not significantly different from that of the dams in three of the four litters when pairwise PERMANOVA was applied to theta distances across age groups (**Fig 2A, Table S1A**). Clustering all the samples together revealed a similar pattern but with distinctions in community structure between different litters (**Fig 2B**). We next compared the similarity between neonates of different ages to the community of their dam versus other dams. Similarity to the dam was lowest at early time points and rose as pups aged to P15 (**Fig 2C**). Samples from pups collected after P15 showed significantly more similarity to their dam than to other dams, whereas early-life samples did not exhibit this pattern (**Fig 2C**). These observations were also evident when samples were clustered based on the Jaccard similarity index, which only considers the presence or absence of OTUs (**Fig S5A-C, Table S1B**). These data demonstrate that pups begin to acquire an adult-like bacterial community beginning around P15, after which the structure of their microbiota coalesces towards their dam’s fecal microbiota.

**Figure 2:**
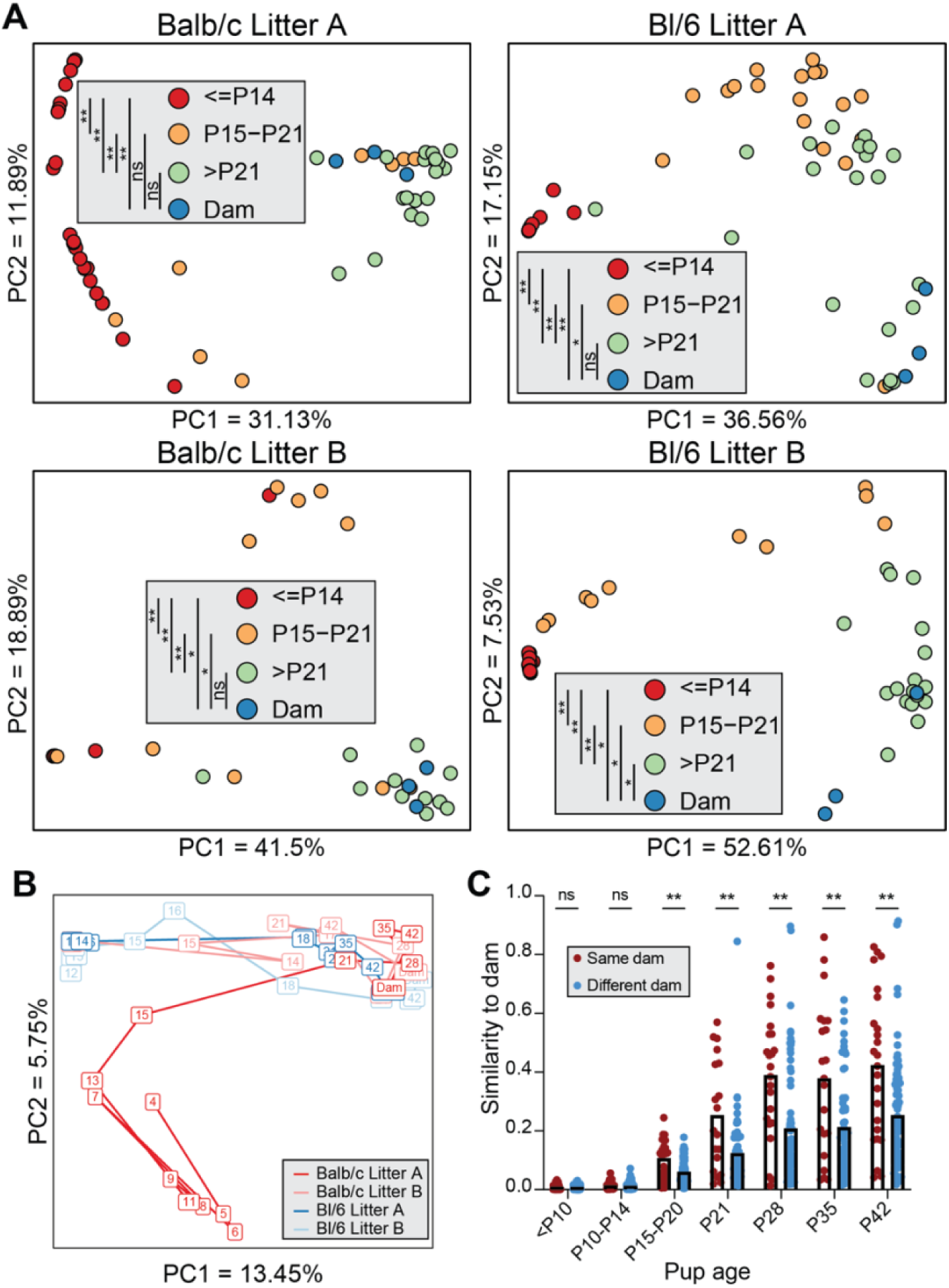
Pup fecal bacterial community structure shifts with age and shares features with dam fecal microbiota. **(A)** Stool samples were clustered by litter using principal coordinate analysis (PCoA) based on theta similarity coefficients. Each point represents a single stool sample, colored according to age. Samples clustered together have a more similar community structure. **(B)** All stool samples were clustered using PCoA based on theta similarity coefficients. Each box represents the average of all stool samples taken at a given age for that litter, with the number indicating the postnatal day on which the samples were collected, with lines connecting subsequent times. **(C)** Theta similarity of samples of the indicated pup age compared to dam samples collected at the age of pup weaning. Samples from each litter were compared either to their dam or to other dams. Means are indicated by the top of the bars. Results were compared by the Kruskal-Wallis test. \**\*\* p* < 0.001, \**\*\*\* p* < 0.0001, ns = not significant; *n* =19-168 pup-dam pairs per group.

### Maternal body sites harbor distinct microbial populations from fecal samples

As the early-life microbiota was quite distinct from the dam fecal microbiota, we asked whether the neonatal microbiome was sourced from other body sites of the dam, as has been seen for humans [1, 7]. To characterize the composition of these bacterial communities, we collected skin swabs (face and ventral) and oral and vaginal washes from the dam shortly after birth and again at pup weaning and sequenced the 16S rRNA V4 gene region, as well as negative controls. While most of the low biomass body site samples exhibited greater read-depth than controls, some samples were excluded from further analysis based on read-depth below 1500 (**Fig S1**). Sequencing of negative control samples revealed “kitome” contaminants, including Pseudomonadaceae, Moraxellaceae, and Comamonadaceae, among others [39] (**Fig S6**). Taxonomic classification of maternal samples showed that samples were generally dominated by Firmicutes, with skin sites having a high representation of Staphylococcaceae, Lachnospiraceae, and Streptococcaceae, oral samples abundant in Staphylococcaceae and Streptococcaceae, and vaginal samples dominated by Staphylococcaceae, Morganellaceae (*Proteus*), or mixed populations (**Fig 3A**). Oral and vaginal samples were significantly less diverse than maternal stool samples (*p*=0.0037 and *p*=0.0312, respectively) (**Fig 3B**). Analysis of bacterial community structure by PCoA of theta similarity showed that fecal samples generally clustered separately from other maternal body sites, but the remaining sites did not cluster by sample type (**Fig 3C**). Overall, this data indicates that there may be substantial overlap in the taxonomic composition of skin, oral, and vaginal taxa of SPF mice, but that these are taxonomically distinct from the enteric microbiome.

**Figure 3:**
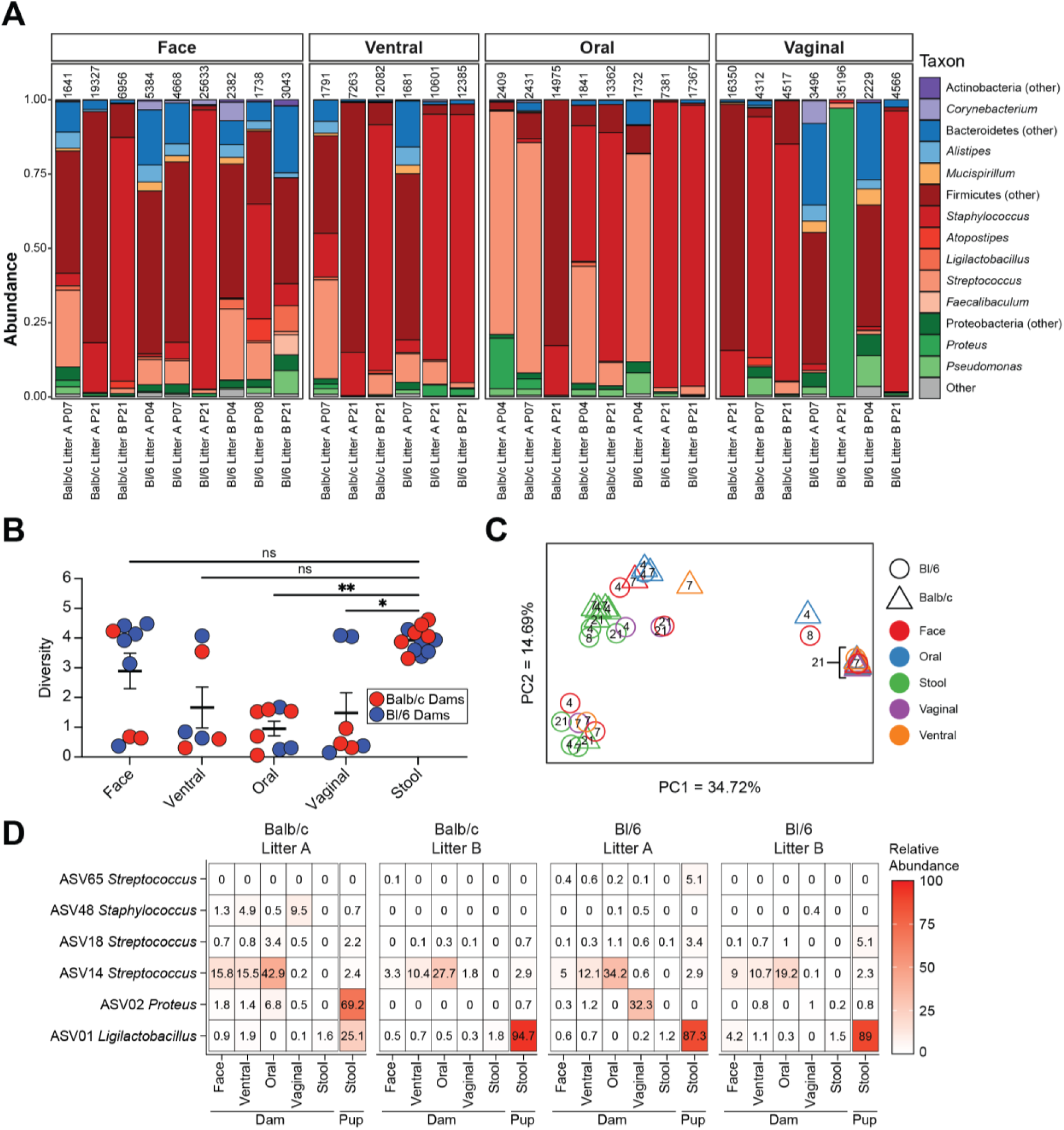
Maternal body site microbiota samples do not cluster by site. **(A)** Taxonomic classification of maternal body-site samples. Genera represented at greater than 5% abundance in at least one sample are shown. Read counts are displayed above each sample; only maternal samples with greater than 1500 reads were used for analysis. **(B)** Shannon diversity of maternal body-site samples. Means are indicated by the thick horizontal crossbar and error bars indicate the standard error of the mean. Results were compared by the Kruskal-Wallis test with Dunn’s test for multiple comparisons. * *p* < 0.05, *\*\* p* < 0.01, ns = not significant; *n* = 6 – 12, representing samples from mice combined across four litters from two genotypes. **(C)** Maternal body site samples clustered using principal coordinate analysis based on theta similarity coefficients. Numbers within points indicate the age of pups at the time of sampling. **(D)** Heatmap of mean relative abundance at maternal body sites of the six ASVs present at greater than 5% relative abundance in the stool of at least one pup up to P10. n = 2-20 samples.

### Dominant early-life taxa are rare but present, in maternal samples

To identify early-life microbes which may be sourced from maternal body sites, we identified ASVs prevalent in pups younger than P10. Across all litters, six ASVs were dominant in early life – one classified as *Ligilactobacillus* (ASV01), one as *Proteus* (ASV02), one as *Staphylococcus* (ASV48), and three as *Streptococcus* (ASV14, ASV18, ASV65). We assessed the abundance of these ASVs in all maternal samples (**Fig 3D**, **S7**). ASV01 and ASV02, the most prevalent in pup fecal samples, were generally rare in all maternal body sites. ASV14, present at lower levels in pup samples, was prevalent in maternal skin and oral samples. Based upon the low-level presence of these ASVs in all maternal sites sampled, a clear maternal source for the pioneering microbes that predominate in early-life fecal samples did not emerge.

### Limited strain-level variance detected in the early pup microbiome

To further resolve the pioneering microbes of the murine neonatal microbiota, we performed shotgun metagenomic sequencing on samples enriched for dominant *Ligilactobacillus* or *Proteus* OTUs (**Table S2,** sequenced samples labeled with dots in **Figure S2**). Metagenome-Assembled Genomes (MAGs) were generated using MEGAHIT [30] and binned using anvi’o [31] (**Table S3**). Taxonomy was assigned to reads using Centrifuge [32], and then reads were mapped to contigs using bowtie [32], permitting taxonomy assignment to contigs and bins. Three large bins were generated corresponding to *Proteus mirabilis* [100% completion, 0% redundancy], *Streptococcus haloterans* [97.2% completion, 4.2% redundancy], and *Ligilactobacillus murinus* [98.6% completion, 1.4% redundancy], validating genus and species assignments generated from the 16S rRNA gene V4 sequencing. Taxonomic assignments for remaining bins included contigs assigned to *Enterococcus*, *Leuconostoc*, *Lactobacillus*, *Lactococcus*, and multiple *Streptococcus* species, eukaryotes, viruses, and mobile genetic elements. Bacterial 16S rRNA gene sequences were recovered from three MAGs – the *Ligilactobacillus murinus* bin contained a match to ASV01, the *Proteus mirabilis* bin matched ASV02, and a MAG predicted to be from *Streptococcus danieliae* contained a match to ASV18 (**Table S3)**.

Strain level diversity was characterized using StrainGST and StrainGR within the StrainGE analysis toolkit (v 1.3.3) [40]. All complete NCBI genomes within *Streptococcus*, *Proteus*, *Ligilactobacillus, Enterococcus*, *Leuconostoc*, *Lactobacillus*, and *Lactococcus* were downloaded and used to build a StrainGST database [41]. Within this subset of samples, these were the top genera identified by 16S rRNA V4 or shotgun metagenomic sequencing and had multiple sequences assigned to them in the previously described analyses. *Ligilactobacillus murinus* ASF361 was the reference strain identified as the best hit across all samples (**Fig 4A**; ∼99.94% nucleotide similarity to reference). Comparison of strains between samples using StrainGR found that when detected there is no significant difference in the genomes of *Ligilactobacillus murinus* collected from Bl/6 or BALB/c mice, or from different mice in different litters (**Fig 4B)**. There were two strains of *Proteus mirabilis* (N18-00201 and swupm1) that were identified as being highly similar when reads were searched against the StrainGST database. These strains were both identified within individual samples and were collapsed to just *Proteus mirabilis* swupm1 as the best representative (**Fig 4B**; ∼99.62% nucleotide identity). This suggested that there was a single strain of *Proteus mirabilis* across all samples tested that shared a high level of similarity with both the N18-00201 and swupm1 reference genomes. There was no significant difference in the genomes of *Proteus mirabilis* collected across mouse lines or litters (**Fig 4C)**. Although a MAG was generated for *Streptococcus haloterans*, no confident strain assignments were generated for any *Streptococcus,* which was present at low relative abundance across all samples and showed evidence of multiple species being present in the MAG data.

**Figure 4:**
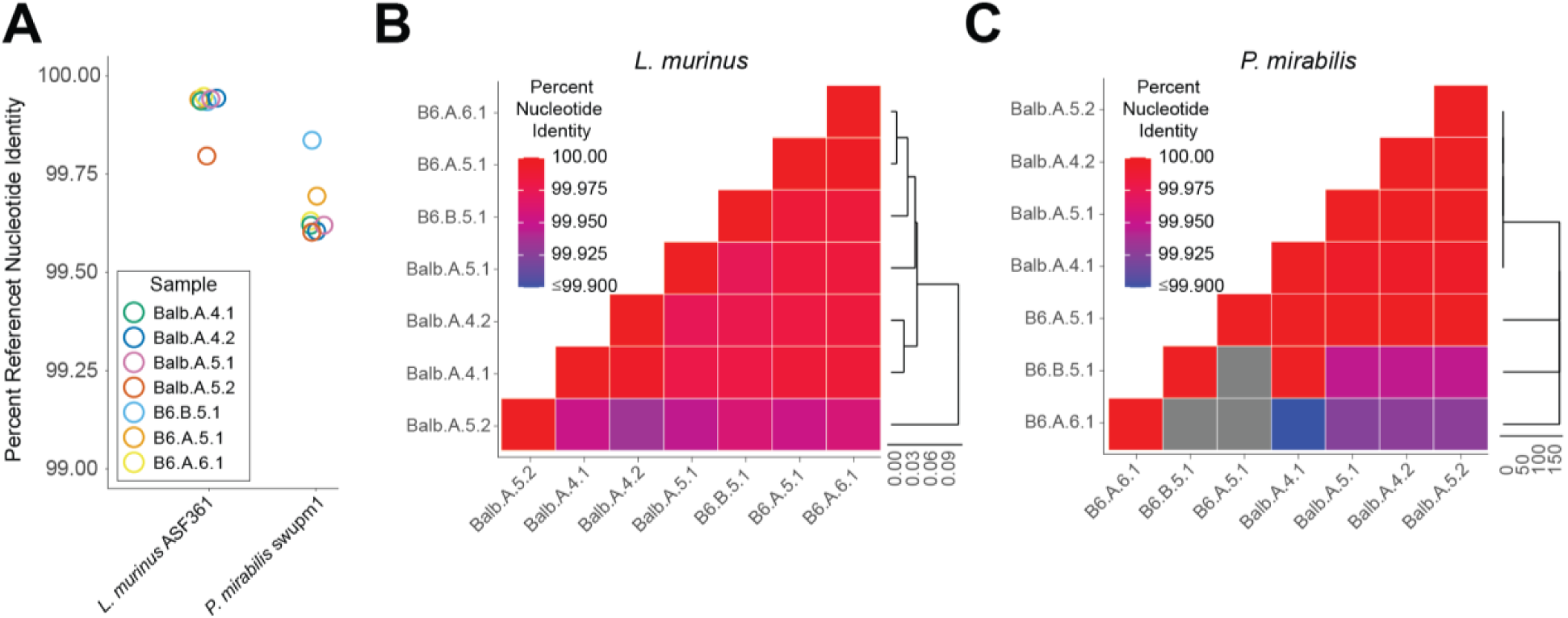
Limited strain-level diversity detected in the early pup microbiome. **(A)** Percent of nucleotide identity shared between strains of *Ligilactobacillus murinus* and *Proteus mirabilis* identified in seven shotgun-sequenced mouse samples and their most closely related NCBI genomes, *L. murinus* ASF361 and *Proteus mirabilis* swupm1, respectively, as determined by StrainGST. Sample names indicate mouse strain, litter, pup age in days, and sample number, separated by dots. **(B)** Heatmap colors indicate pairwise nucleotide identity between *L. murinus* strains identified in each of the seven mouse samples as determined by StrainGR. The cladogram was calculated using Euclidean distance. **(C)** Heatmap colors indicate pairwise nucleotide identity between *P. mirabilis* strains identified in each of the seven mouse samples as determined by StrainGR. The cladogram was calculated using Euclidean distance.

### Viruses and plasmids identified in the early pup microbiome

Although minimal variation in bacterial strains was observed across individual mice, mouse strains, or litters, we sought to evaluate other potential sources of genomic variation by examining flexible genomic regions including viruses and plasmids. During binning, one bin was generated containing a single contig that was an exact match to a known *Streptococcus* phage. We next performed a detailed search for viruses present in the early stages of murine microbiome development, identifying 31 viral-like regions across four bacterial MAGs using VirSorter2 [42](**Table S4**). Predicted viral regions covered from 0.14% to 5.55% of the length of these four MAGs (**Table S3**). While predicted viruses ranged in length from 1,094 bp to 94,000 bp, with 26 covering over 80% of the length of the contig they were identified in, they tended to be short with a median length of 3,307 bp and 21 < 5,000 bp. VirSorter2 identified 24 as dsDNA phages and 7 as ssDNA viruses. We next sought to characterize this collection of viruses, while recognizing that their short, fragmented, nature may hinder bioinformatic predictions. Ten of these viruses were predicted to be temperate, while six were predicted to be virulent, and the remainder were filtered or did not receive a lifestyle prediction from PhaTYP [33] (**Table S4**). Five viral regions received confident taxonomic predictions from PhaGCN and were assigned to four viral families: *Drexlerviridae* (2), *Peduoviridae* (1), *Casjensviridae* (1), and *Straboviridae* (1) [34]. We subsequently used mmseqs2 to search a database of non-redundant nucleotide sequences compiled from the gut phage database (GPD) and identified 22 phages with E-value ≤ 4.09 x 10^-4^ [35, 36]. Seven additional viral sequences from our dataset were assigned putative taxonomic predictions as *Siphoviridae* (4), *Myoviridae* (2), and *Podoviridae* (1) based on the metadata associated with their top hit in the GPD. The thirteen GPD representatives with associated host range information generally matched the taxonomy of the MAG where the virus was binned (11 matched genus, 1 matched family, and 1 matched order). The median abundance of viral contigs (transcripts per million) was not significantly elevated relative to other contigs in the bin within any of the samples (Mann–Whitney U-test followed by Bonferroni correction) (**Table S4**). Similarly, the number of reads mapping within integrated prophages, defined as viral regions that covered less than 80% of their assembled contig (minimum contig length of 3,000bp), were not significantly different than those mapping to the contig outside of the prophage region. This suggests that the phages identified by VirSorter occur with similar copy numbers to their bacterial host genome, suggesting they are lysogenic.

During binning with anvi’o, four contigs were predicted to be plasmids from *Enterococcus* (1 contig), *Proteus* (2 contigs), or *Staphylococcus* (1 contig) based on their similarity to plasmid sequences in the NCBI nucleotide database and differential clustering compared to the dominant bacterial bins (**Table S3**). This encouraged the bioinformatic prediction of plasmid sequences in our dataset that may not have been binned separately from their host genomes due to limited variation in sequence composition and coverage across samples. We applied Plasmer [37] to our assembled contigs and identified 25 plasmid-like contigs that ranged in length from 1,013 bp to 8,057 bp, sixteen of which were found in non-mammalian bins (**Table S5**). Plasmer not only confirmed that the four contigs identified during binning originated from plasmids, but also resolved their predicted taxonomic origin (*Enterococcus faecalis*, *Proteus mirabilis*, and *Staphylococcus aureus*). Plasmer additionally identified plasmids that were predicted to originate from *Ligilactobacillus murinus* and *Streptococcus*, suggesting that predominant taxa of early microbiome development have associated plasmids.

## Discussion

In this work, we closely characterized the development of the murine gut microbiota over the first weeks of life. The early-life microbiota is extremely simple, consisting of primarily *Ligilactobacillus*, *Proteus*, and *Streptococcus*. These taxa are rare in maternal fecal samples and body sites, so selection for these taxa likely occurs in the neonatal gut after exposure to these microbes from the dam or other untested sites such as breastmilk or bedding. Around P15, the gut microbiota begins to increase in diversity and shifts dramatically in composition, with Bacteroidetes, including *Bacteroides*, as well as a more diverse set of Firmicutes replacing the dominant early-life taxa. By the age of weaning at P21, diversity has stabilized, and the pups have acquired a microbial community that most closely resembles their dam. The in-depth longitudinal characterization of the early-life bacterial microbiota performed here demonstrates the notable shifts during microbiota maturation that occur pre-weaning and provides a framework to explore phenotypes affected by neonatal microbiota.

The simplistic bacterial communities common in neonatal mice are in line with studies demonstrating that human infant gut microbiota is low diversity and becomes more complex over the first years of life [3, 13]. The dominance by *Ligilactobacillus* (a newly described genus, previously included in the *Lactobacillus* genus) [43] in some ways mimics features seen in human infant microbiotas. *Lactobacillus* colonizes vaginally-born infant*s* at many body sites (including skin, oral cavity, and nose), consistent with exposure to the mother’s vaginal microbiota, which is often dominated by Lactobacilli, particularly in pregnant women [44]. *Lactobacillus* is also enriched in breast-fed infants [2, 8, 45], in part because this taxon is present in human breast milk [46]. Additionally, *Lactobacillus* species identified in these previous human studies may have since been reclassified as *Ligilactobacillus* [43]. However, human infants are typically colonized by Bifidobacteria [2, 45], a taxon that was not present in the neonatal mouse samples, consistent with previous studies [47]. As colonization with *Bifidobacterium* species in infancy is thought to play an important role in human health [48], this represents a key distinction between the microbial communities of neonatal mice and humans.

The dramatic increase in diversity and shift in microbial community structure that occurs at P15 in neonatal mice is consistent with the age at which pups shift from a breastmilk-exclusive diet and begin eating solid food [49]. This is a gradual process wherein the proportion of breastmilk in the diet decreases until weaning when the pups shift exclusively to solid food [50] and is accompanied by changes in intestinal gene expression related to the metabolism of dietary macromolecules and immune responses [51–53]. The change in nutrient availability allows for colonization by a diverse set of microbes – however, the diet is not the only selective factor, as a neonatal microbial community can stably colonize germ-free mice even when they are fed a solid food diet [54, 55]. This is also the period in which pups begin to exhibit coprophagy, allowing the transfer of fecal microbes directly from the dam to neonates [56]. This fecal-oral transfer of microbes likely explains why microbial populations in mice most closely resemble their nursing dam [20, 21].

Our analysis did not provide a clear origin of early-life microbes, as the microbes dominant in the neonatal gut were rare at all sites tested. Low levels of microbes from these maternal body sites may seed the neonates, which then expand in the neonatal gut [57]. However, it is also possible that other sources of microbes, such as breastmilk [58], may seed the neonatal gut. Strain-level resolution of the microbes present at these sites could suggest a most likely origin for the neonatal microbiota.

Exposure to the gut microbiota in early life is extremely important for later-life health, and disruption of the neonatal microbiota can lead to long-lasting metabolic and immune dysfunction [59, 60]. This work closely characterizes the development of the murine gut microbiota over the first weeks of life, providing a detailed understanding of the specific bacterial taxa present at different developmental time points in this important model organism, highly relevant for the study of early-life challenges and exposures.

## Conclusions

The murine bacterial microbiota begins as a simple community dominated by a handful of pioneering, milk-associated, taxa. There is not a single source for these bacteria; they are found at low levels at multiple maternal body sites. After 14 days of postnatal life, the gut bacterial community of pups rapidly changes, becoming significantly more diverse and similar to their dam – this process is largely complete by day 21. This developmental process is an important determinant of host health and alterations of this program have been implicated in disease processes. Future studies attempting to modulate this period of maturation will be greatly informed by our detailed analysis of the process across multiple mouse strains and litters.

## List of abbreviations

ASV: Amplicon sequence variant
GPD: Gut phage database
LEfSe: Linear discriminant analysis effect size
MAG: Metagenome-assembled genome
OTU: Operational taxonomic units
PCoA: Principal coordinate analysis
SPF: Specific pathogen-free

## Declarations

### Ethics approval and consent to participate

Not applicable

### Consent for publication

Not applicable

### Availability of data and materials

Data and files necessary to generate figures and statistical analyses, as well as ASV data, metagenomic assemblies, and plasmid and viral predictions, have been uploaded to Zenodo (DOI:10.5281/zenodo.10456555). These include the input data, R markdown files, and Prism files, a record of analyses run generated using the ‘knitr’ package in R [61], and the input data for Prism analyses exported as Excel or text files. Sequencing reads and associated metadata for V4-16S rRNA gene and short-read shotgun metagenomic sequencing have been uploaded to the Sequence Read Archive (BioProject ID: PRJNA1061151).

### Authors’ contributions

E.A.K., A.H.K., A.L.D., and A.A. performed the experiments. E.A.K. and J.S.W. analyzed the data and generated figures. E.A.K., J.S.W., and M.T.B. wrote the paper. All authors read and edited the paper.

## Acknowledgments

Not applicable

