## Supplemental Figures 1-7 for "Microbiota assembly of specific pathogen-free neonatal mice"

Supplemental Data for "**Microbiota assembly of specific pathogen-free neonatal mice**"

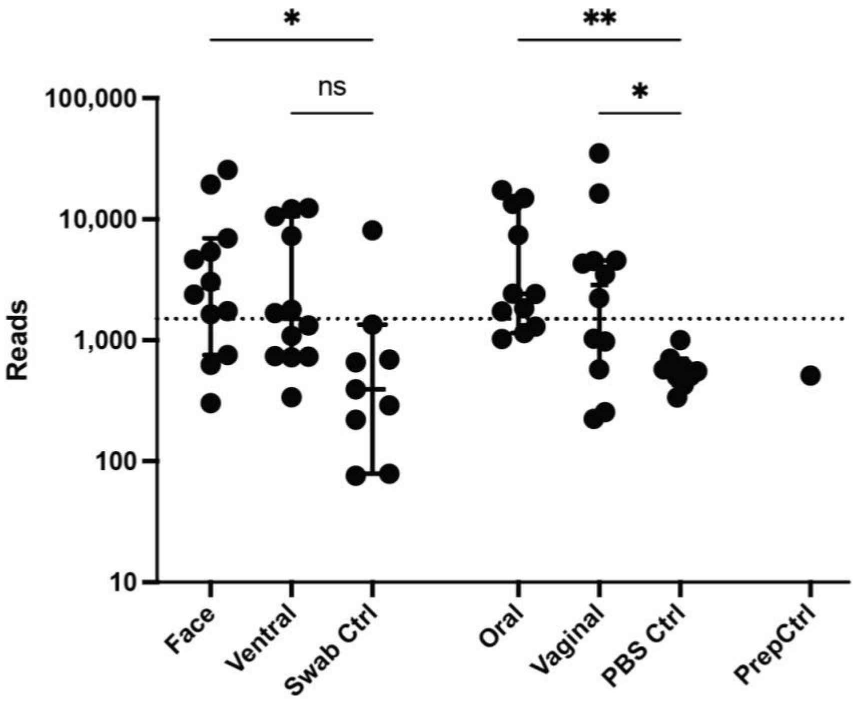


**Figure S1. Sequencing of negative control samples revealed fewer reads than maternal body site samples.**

Post-processing read counts for all samples and controls. PBS and swab samples were collected alongside each set of maternal samples, acting as controls for oral and vaginal washes, and skin samples, respectively. The prep control was a negative control included for sample processing. Error bars represent the median and 95% confidence interval. Kruskal-Wallis followed by Dunn's multiple comparisons test was used to compare the medians of face (n=12) and ventral (n=12) samples to swab-only (n=9) controls and oral (n=11) and vaginal (n=12) wash samples to the PBS (n=9) control. Samples with >1500 reads (dashed line) were included in analyses. * $p < 0.05$, ** $p < 0.01$, ns = not significant.

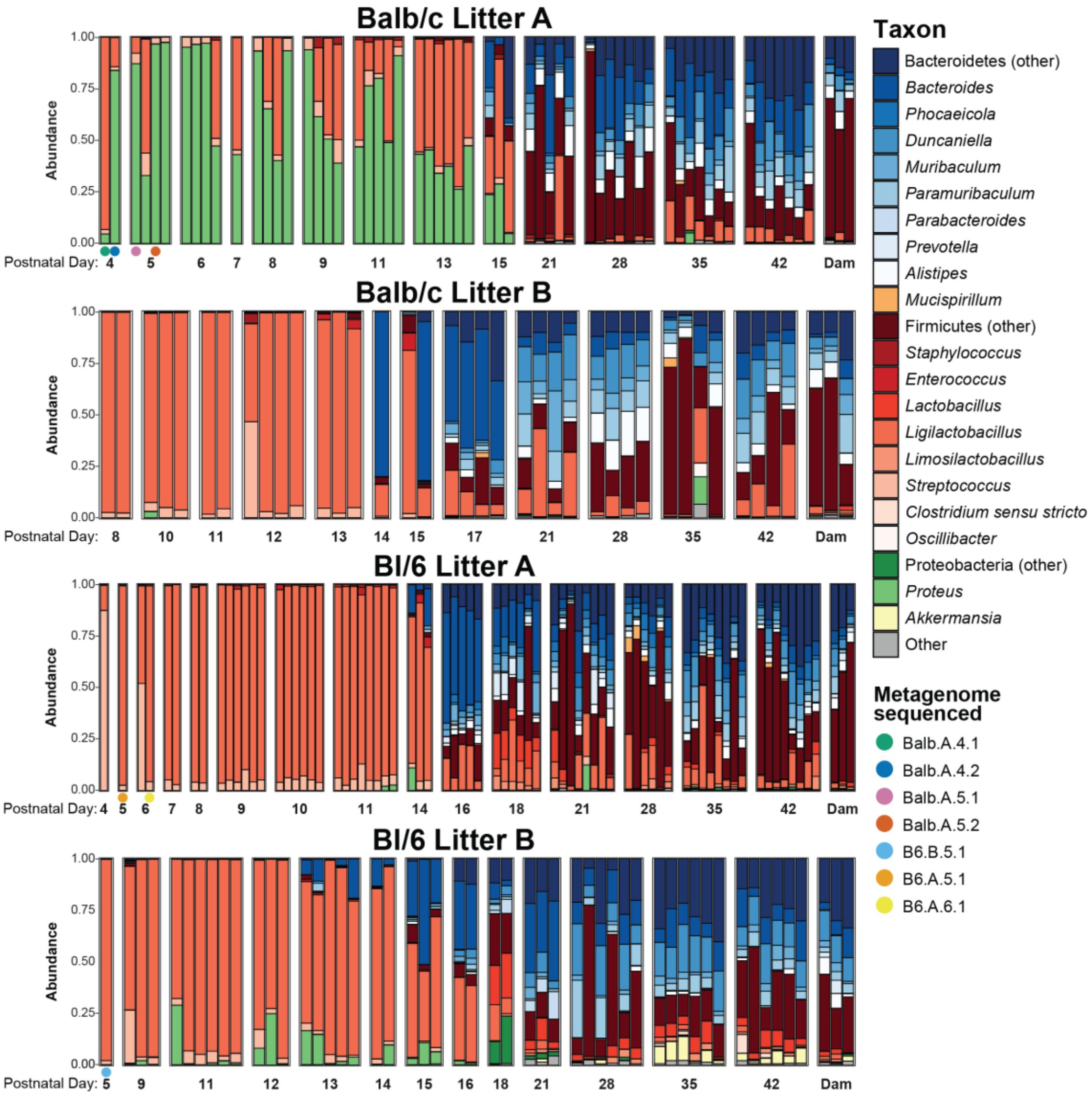


**Figure S2: Taxonomic assignments of longitudinal fecal samples**

Taxonomic classification of stool samples collected from neonates sequenced at the V4 region of the 16S rDNA gene. Genera represented at greater than 5% abundance in at least one sample are shown. Each bar represents a single fecal sample collected at the indicated postnatal day. Color families represent phyla-level assignments – blue for Bacteroidetes, orange for Deferribacteres, red for Firmicutes, green for Proteobacteria, and yellow for Verrucomicrobia; grey includes phyla present at less than 5% abundance in all samples. Colored dots underneath bars indicated samples that were short-read shotgun sequenced.

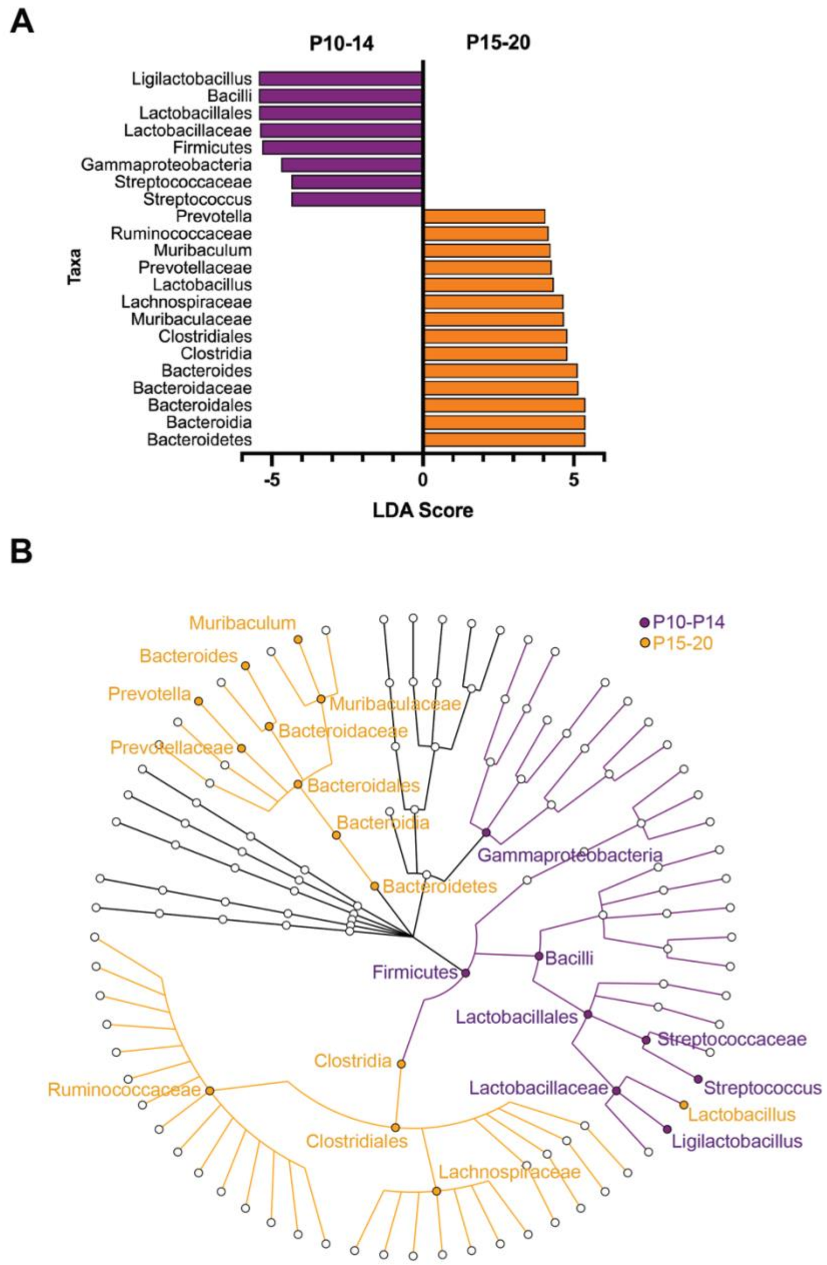


**Figure S3: Differential pre-weaning taxa before and after P14.**

LEfSe analysis was performed to identify discriminatory taxa between pup microbiota samples from P10-P14 versus P15-P20. Cutoff was a logarithmic linear discriminant analysis (LDA) score of 4.0. **(A)** Discriminatory taxa displayed based on effect size. **(B)** Cladogram of age-discriminant taxa.

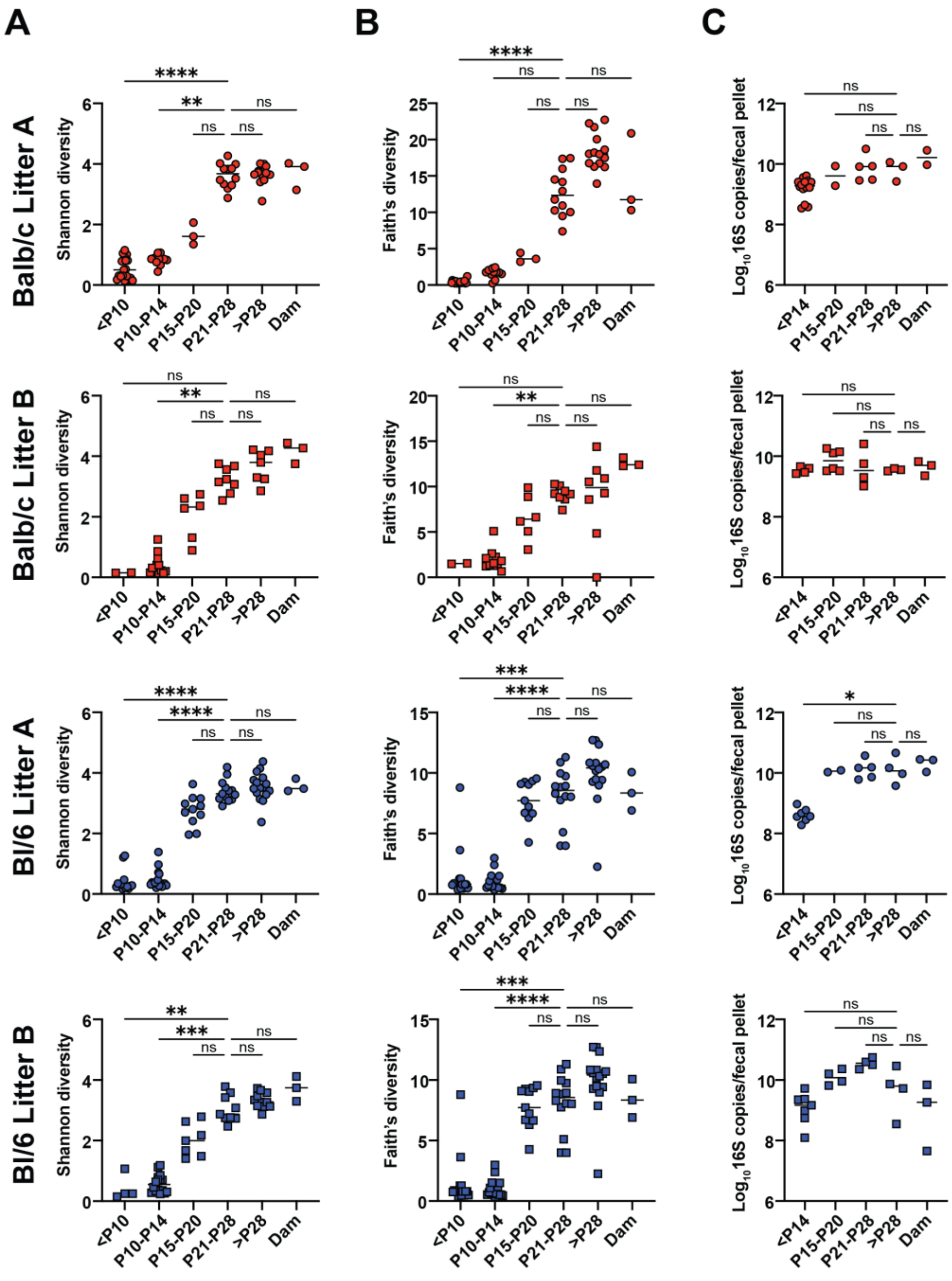

**Figure S4: Alpha diversity analysis by litter reveals consistent longitudinal increases in diversity prior to weaning.**

**(A)** Shannon diversity calculated based on operational taxonomic unit (OTU) clustering. **(B)** Faith's phylogenetic diversity calculated based on phylogeny of amplicon sequence variants (ASVs). **(C)** 16S rDNA copies per fecal pellet, detected by qPCR. Means are indicated by the horizontal lines. Results were compared by the Kruskal-Wallis test. * $p < 0.05$, ** $p < 0.01$, *** $p < 0.001$, **** $p < 0.0001$, ns = not significant; $n$ =2-19 mice per experimental group.

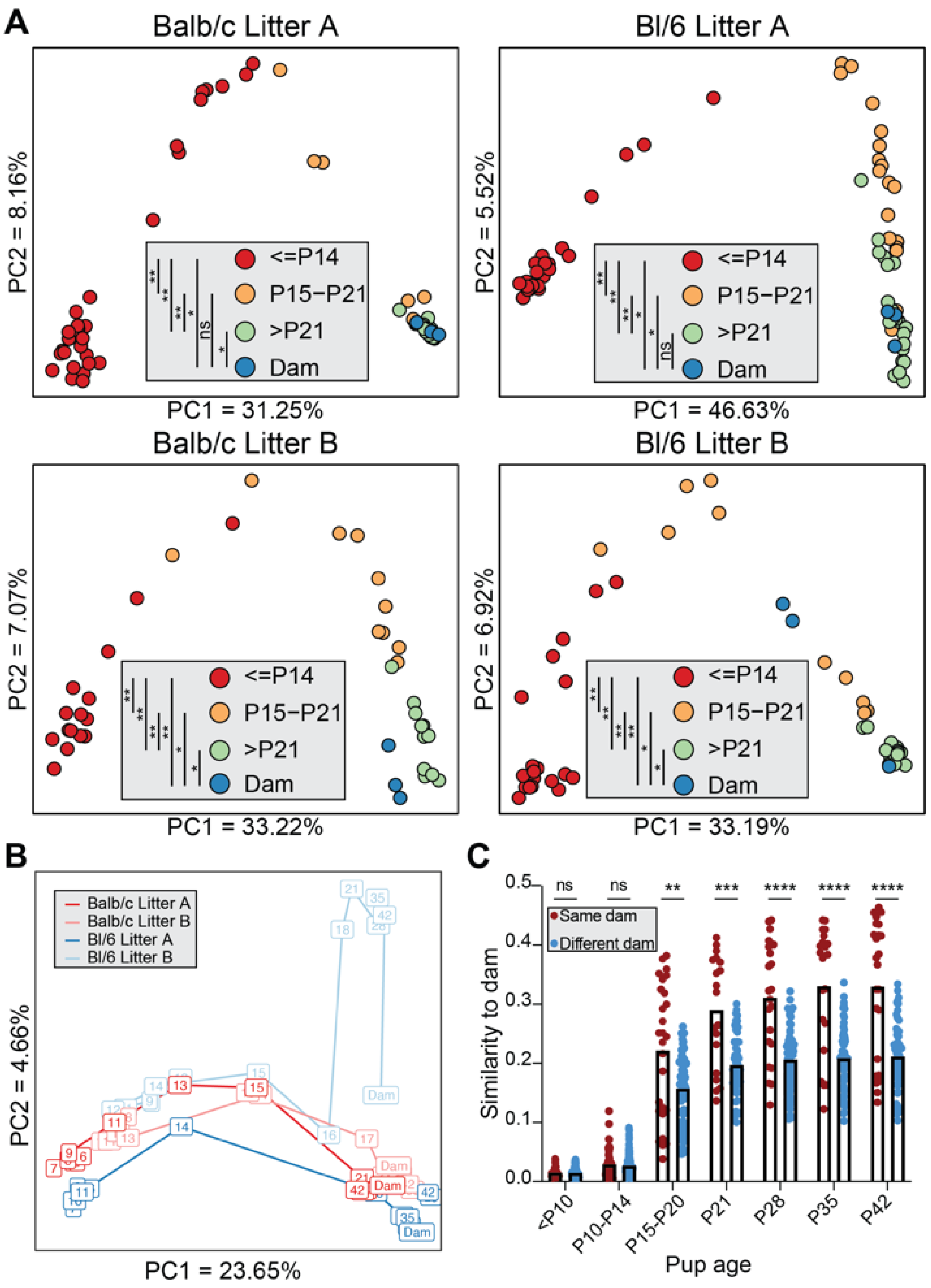


**Figure S5: Jaccard index clustering of fecal samples**

**(A)** Principal coordinate analysis based on Jaccard similarity coefficients was performed on stool samples from each litter. Each point represents a single stool sample, colored according to age. Samples clustered together have more similar community structure. **(B)** Ordination was performed on all stool samples using principal coordinate analysis based on Jaccard similarity coefficients. Each box represents the average of all stool samples taken at a given age for that litter, with the number indicating the postnatal day on which the samples were collected, with lines connecting subsequent times. **(C)** Jaccard similarity of samples of the indicated pup age compared to dam samples collected at the age of pup weaning. Samples from each litter were compared either to their own dam or to other dams. Means are indicated by the top of the bars. Results were compared by the Kruskal-Wallis test. ** $p < 0.01$, *** $p < 0.001$, **** $p < 0.0001$, ns = not significant; $n$ =19-168 pup-dam pairs per group.

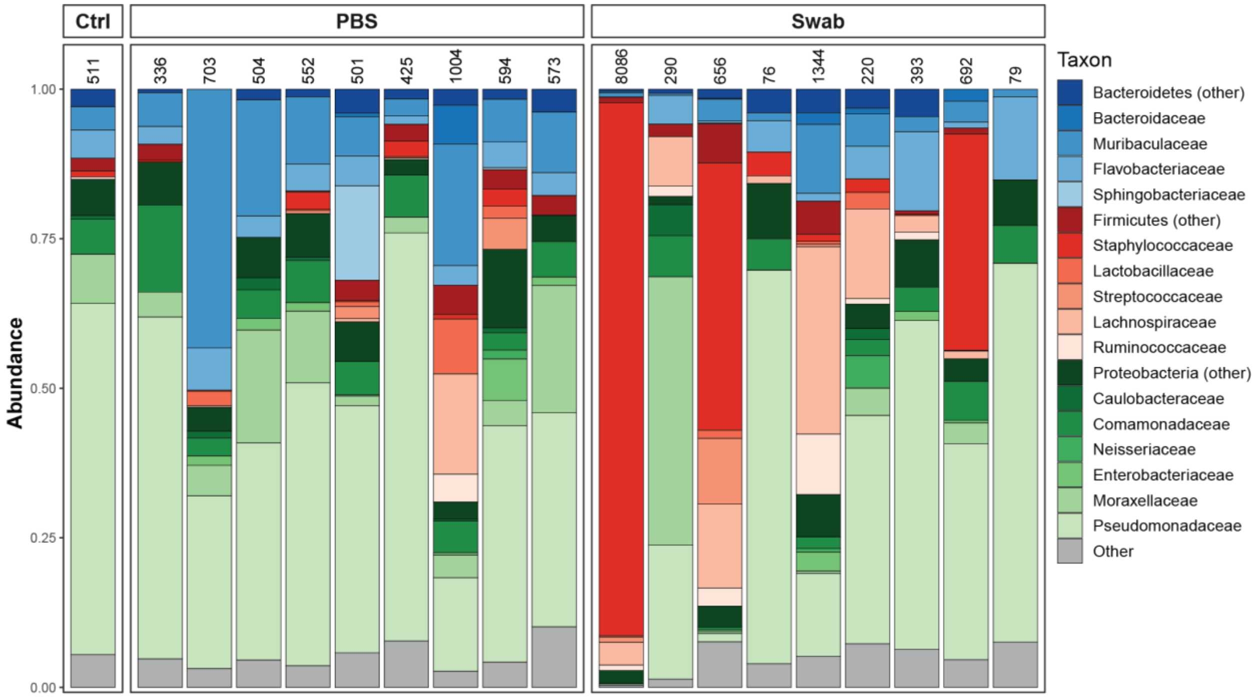


**Figure S6: Sequencing of negative control samples revealed presence of common "kitome" contaminants.**

Taxonomic classification of negative controls; families represented at greater than 5% abundance in at least one sample are shown. Read counts are displayed above each sample.

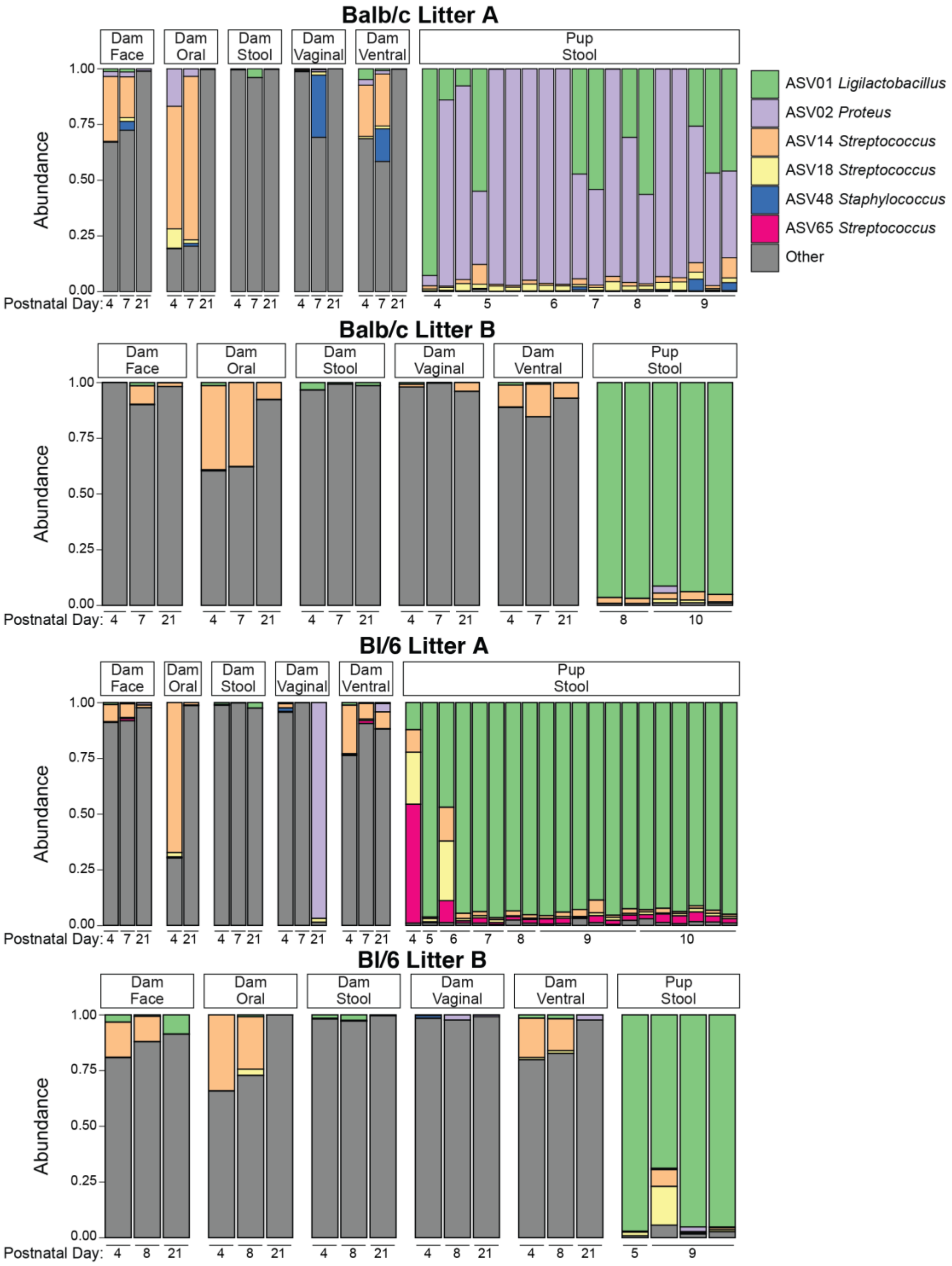


**Figure S7: Dominant early-life taxa are rare in maternal samples.**

ASVs present at greater than 5% in any pup sample collected up to P10 were quantified in maternal samples and neonatal stool samples.
